# Replication efficiencies of human cytomegalovirus-infected epithelial cells are dependent on source of virus production

**DOI:** 10.1101/2024.03.19.585739

**Authors:** Rebekah L. Mokry, Christopher E. Monti, Suzette Rosas-Rogers, Megan L. Schumacher, Ranjan K. Dash, Scott S. Terhune

## Abstract

Human cytomegalovirus (HCMV) is a prevalent betaherpesvirus, and infection can lead to a range of symptomatology from mononucleosis to sepsis in immunocompromised individuals. HCMV is also the leading viral cause of congenital birth defects. Lytic replication is supported by many cell types with different kinetics and efficiencies leading to a plethora of pathologies. The goal of these studies was to elucidate HCMV replication efficiencies for viruses produced on different cell types upon infection of epithelial cells by combining experimental approaches with data-driven computational modeling. HCMV was generated from a common genetic background of TB40-BAC4, propagated on fibroblasts (TB40_Fb_) or epithelial cells (TB40_Epi_), and used to infect epithelial cells. We quantified cell-associated viral genomes (vDNA), protein levels (pUL44, pp28), and cell-free titers over time for each virus at different multiplicities of infection. We combined experimental quantification with data-driven simulations and determined that parameters describing vDNA synthesis were similar between sources. We found that pUL44 accumulation was higher in TB40_Fb_ than TB40_Epi_. In contrast, pp28 accumulation was higher in TB40_Epi_ which coincided with a significant increase in titer for TB40_Epi_ over TB40_Fb_. These differences were most evident during live-cell imaging, which revealed syncytia-like formation during infection by TB40_Epi_. Simulations of the late lytic replication cycle yielded a larger synthesis constant for pp28 in TB40_Epi_ along with increase in virus output despite similar rates of genome synthesis. By combining experimental and computational modeling approaches, our studies demonstrate that the cellular source of propagated virus impacts viral replication efficiency in target cell types.

**IMPORTANCE:** Human cytomegalovirus (HCMV) is a ubiquitous pathogen that can cause serious disease under conditions of immunodeficiency and upon congenital infection. HCMV replicates in diverse cell types throughout the human body with tropism influenced by the source of the virus. Here, we investigated the contribution of viral sources to the kinetics of HCMV replication in epithelial cells using both experimental and mechanistic computational modeling approaches. These studies reveal that HCMV produced from epithelial cells exhibits a higher efficiency of replication despite similar viral DNA synthesis kinetics between viral sources. These differences likely involve a propensity of epithelial-derived virus to induce syncytia versus fibroblast-derived virus, and an accompanying higher synthesis rate of a late virion protein ultimately resulting in production of more extracellular infectious virus.

## INTRODUCTION

Human cytomegalovirus (HCMV) is a betaherpesvirus with an estimated global seropositivity rate of approximately 83% (1). HCMV is the leading cause of congenital birth defects (2, 3) and a major cause of morbidity and mortality in immunocompromised individuals such as hematopoietic stem cell or solid organ transplant patients (4, 5). Infection causes a range of diseases in immunocompromised people including CMV retinitis, pneumonitis, hepatitis, and gastrointestinal disease (4, 6). HCMV pathogenicity is likely facilitated by the ability of the virus to successfully infect and replicate in various tissues and cell types in the body. Disease onset is initiated by primary infection or latency reactivation events that result in lytic replication and dissemination of the virus into different cell types and tissues (7, 8). Although several effective antivirals are available, HCMV infection continues to be a major challenge in healthcare across the globe.

HCMV lytic replication cycle defined using fibroblast cells is marked by viral genome synthesis, the temporal expression of viral RNAs and proteins, and the production of new virus (9, 10). Ribosomal profiling studies identified approximately 700 RNAs during lytic replication (11). Kinetics of gene expression have been defined as immediate early (IE), early (E), early-late (E-L) and late (L) expressed genes based on specific experimental criteria (9). More recently, this has been updated using advanced transcriptomics approaches to define temporal RNA classes TC1-7 (12). The exact numbers of viral proteins and proteoforms (i.e., one protein with varying combinations of modifications) remain unknown (13). Of known proteins, these have characteristic temporal profiles (Tp1-5) during single-step replication irrespective of their varying post-translational modifications (10).

To better understand the complex relationships occurring during infection, we and others have begun introducing data-driven mechanistic computational modeling that simulate relationships and changes over time *in silico* (14–16). We recently used the dynamic profiles of two viral proteins to construct a computational model for single-step replication, revealing MRC-5 fibroblast cells have a maximal capacity to produce virus along with an optimal input of HCMV genomes (16). In fibroblasts, the lytic replication cycle has an approximately 96 hour duration that culminates in production of infectious virions and destruction of the infected cell (17). However, replication differences exist between cell types and likely involve differences in inoculum compositions and diverse host cell factors (18–22). These virion and cell-type dependent relationships and associated kinetics remain poorly defined.

In this work, we used experimental approaches complemented by mechanistic computational modeling to investigate differences in HCMV replication efficiencies between virus source and target cells. We define “virus source” as the cell type in which the virus was propagated, while “target cell” is the identity of the uninfected cells that are infected with virus from a specific viral source. We used virus from a common genetic background that was propagated from either fibroblasts or epithelial cells and infected epithelial cells using several MOIs. We quantified numerous features of HCMV replication over time including viral genome synthesis, viral protein levels, and virus production which were used to inform a mechanistic computational model (16). By combining experimental data with mechanistic computational modeling, we have uncovered unique aspects of virus source and target cell type-dependent HCMV replication.

## RESULTS

### Kinetics of viral DNA replication during infection by epithelial- and fibroblast-derived HCMV

Tropism and entry of HCMV is influenced by the cell type in which the virus is propagated (23). We asked whether the virus source from the same genetic background might influence the kinetics of replication on a common target cell, ARPE19 epithelial cells. Viral stocks were prepared by electroporating HCMV TB40/E, clone TB40-BAC4 (24) into MRC-5 fibroblasts (**Fig. 1A**) using a recombinant genome that expresses IE2-T2A-eGFP and pp28-mCherry provided by Eain Murphy (SUNY Upstate Medical University, Syracuse, NY) (16). The resulting viral stock was subsequently used to infect MRC-5 fibroblast cells resulting in an HCMV stock referred to as TB40-BAC4_Fb_ (TB40_Fb_). Alternatively, the initial stock was passaged three times through ARPE-19 epithelial cells resulting in a HCMV stock, TB40-BAC4_Epi_ (TB40_Epi_). Stocks were prepared using cell-free virus and concentrating by sorbitol pelleting. Titers were determined using TCID_50_ assays on the same cell type in which the virus was propagated. We infected ARPE19 epithelial cells at MOIs 0.1 IU/cell and 0.5 IU/cell with either TB40_Fb_ or TB40_Epi_ (**Fig. 1A**). We quantified levels of viral DNA (vDNA) and have included data from Monti et al. (16) investigating TB40_Fb_ infection of MRC-5 fibroblasts (**Fig. 1B**) as a reference point. We isolated cell-associated DNA and measured vDNA by qPCR from 2-96 hpi using primers targeted to the UL123 gene and cellular DNA targeting the CDKN1A gene. We included plasmid standards for each gene resulting in gene copies per microliter of sample for TB40_Fb_ (**Fig. 1C**) and TB40_Epi_ (**Fig. 1D**). For these studies, we refer to the initial 2 hpi time as vDNA_in,0_ (16), representing the amount of cell-associated vDNA at 2 hpi. Viral genomes per cell were measured by normalizing UL123 genome copies (**Figs. 1C, D**) to two CDKN1A genome copies (**Fig. 1E**). The mean vDNA_in,0_ for TB40_Fb_ infection of epithelial cells were 0.3 and 1 genomes/cell for MOIs 0.1 and 0.5 IU/cell, respectively. TB40_Epi_ infection of epithelial cells resulted in 0.6 and 3 genomes/cell for MOIs 0.1 and 0.5, respectively. By 24 hpi, there were no differences in viral genomes levels between TB40_Fb_ and TB40_Epi_ at a given MOI (**Fig. 1E**), suggesting that the intrinsic differences in viral stocks are most evident between 2 and 24 hpi. Despite using the same MOIs, vDNA_in,0_ from our previous studies of TB40_Fb_ infection of fibroblast cells (16) are substantially higher compared to infection of epithelial cells suggesting that use of vDNA_in,0_ is a more consistent measurement of HCMV input. At 96 hpi, we measured similar levels of viral genomes per epithelial cell between TB40_Epi_ and TB40_Fb_ at a given MOI (**Fig. 1E**).

**Figure 1.**
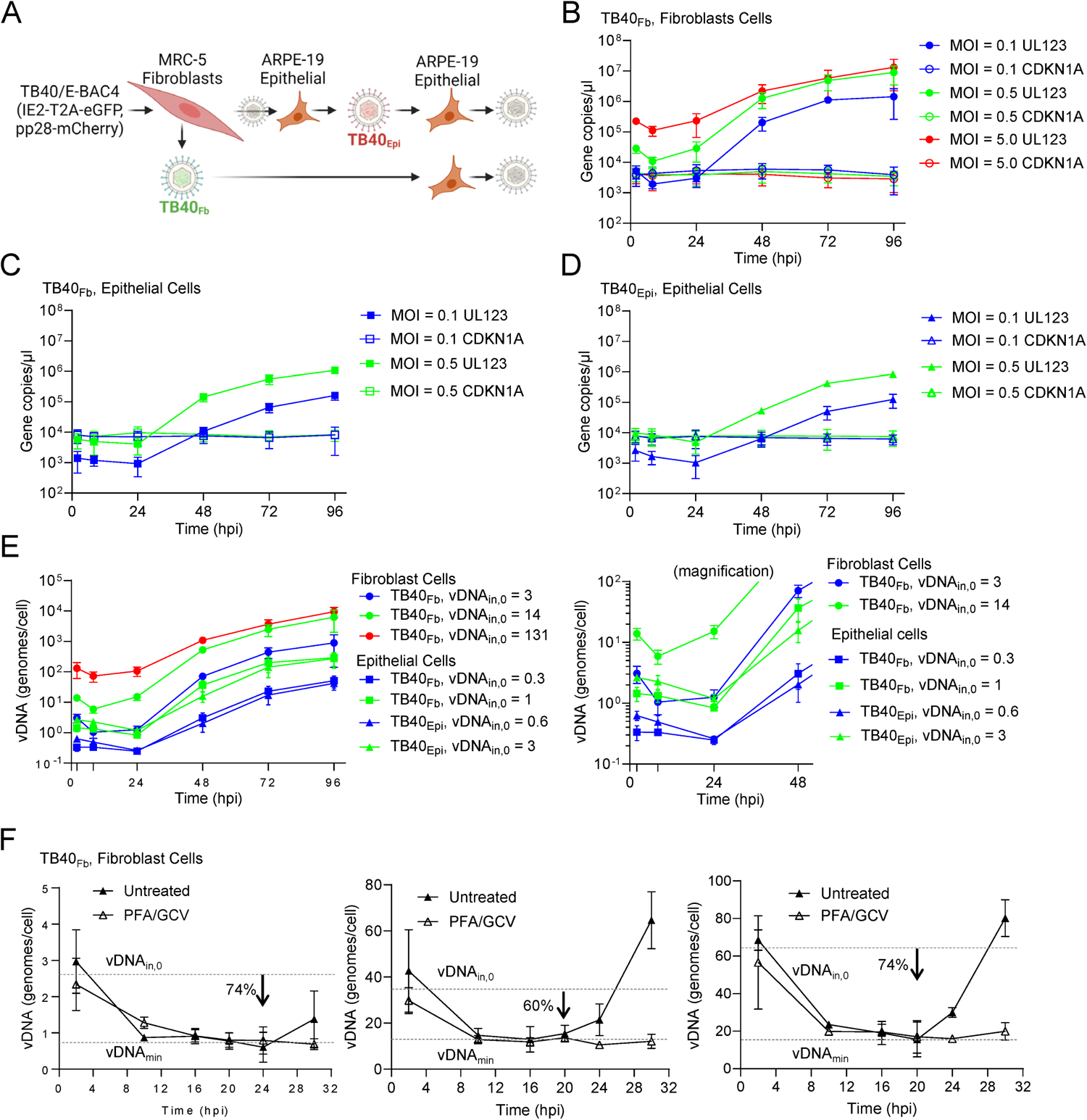
Comparative HCMV genome replication kinetics between sources and target cells. **(A)** Schematic of BAC-derived virus propagation. HCMV TB40-BAC4 expressing IE2-T2A-eGFP and pp28-mCherry was transfected into MRC-5 fibroblasts to produce an initial viral stock. Subsequently, virus stocks for experiments were produced from MRC-5 fibroblasts, TB40_Fb_ or from ARPE-19 epithelial cells, TB40_Epi_. Kinetics of replication were measured by infecting ARPE-19 cells. **(B)** qPCR data from (16) reproduced as comparison control. Confluent ARPE-19 cells were infected at MOI of 0.1 or 0.5 IU/cell using **(C)** TB40_Fb_ or **(D)** TB40_Epi_ viruses. Viral DNA (vDNA) and cellular DNA were quantified using absolute qPCR with primers to viral UL123, cellular CDKN1A, and plasmid standards. Data was obtained from three biological replicate experiments and standard deviations are shown. **(E)** Data are presented as vDNA per cell for each condition. vDNA_in,0_ at 2 hpi for TB40_Epi_ are 0.6 vDNA/cell (MOI 0.1) and 2.6 vDNA/cell (MOI 0.5). TB40_Fb_ are 0.3 vDNA/cell (MOI 0.1) and 1.4 vDNA/cell (MOI 0.5). Right panel is magnified for 0 to 48 hpi. Data were obtained from three biological replicate experiments and standard deviations are shown. **(F)** Confluent MRC5 cells were infected at indicated vDNA_in,0_ with TB40_Fb_. Cells were untreated or treated with 10 mM ganciclovir (GCV) and 30 mg/mL phosphonoformate (PFA). Absolute qPCR data were obtained as described above to define the timing of vDNA_min_ from two biological replicate experiments with standard deviations shown.

Noting the variability in input and initial decrease in viral genomes from 2-24 hpi, we identified the minimum in vDNA levels at different vDNA inputs. Since we are fitting models to these experimental data, vDNA time course measurements using the time points of 2, 8, and 24 hpi might miss these low points and artificially introduce a minimum at an inappropriate time point. Therefore, we increased the resolution of our measurements by increasing the number of time points over a shorter duration to define the minima at three vDNA_in,0_ of approximately 2.7, 36.2, and 62.6 genomes/cell (**Fig. 1F**). We defined the minimum point sustained over several time points while blocking viral DNA synthesis. We quantified changes in vDNA over time of TB40_Fb_ infecting fibroblast cells in the presence and absence of vDNA synthesis inhibitors phosphonoformate (PFA) and ganciclovir (GCV) (**Fig. 1F**). Using this approach, we defined vDNA_in,0_, vDNA_min_ (minimal level), and t_rep_ which is the approximate time of replication onset. We defined the minima for vDNA_in,0_ of 2.7 genomes/cell occurring between 16-24 hpi with t_rep_ at approximately 24 hpi. For vDNA_in,0_ of 36.2 and 62.6 genomes/cell, vDNA_min_ occurred between 10-20 hpi with t_rep_ of 20 hpi. These observations demonstrate that vDNA levels decline to a sustained minimal level (vDNA_min_) prior to the onset of synthesis. vDNA_min_ were found to be approximately 0.7 genomes/cell (vDNA_in,0_ = 2.7), 14.5 genomes/cell (vDNA_in,0_ = 36.2), and 16.3 genomes/cell (vDNA_in,0_ = 62.6) which represents a 60-74% reduction from vDNA_in,0_. Importantly, these results support our previously developed empiric model which inherently involves a minimum around 18-24 hpi (16). Based on our new data, vDNA_min_ is likely within same general window for TB40_Fb_ and TB40_Epi_ infecting epithelial cells (**Fig. 1E**). Applying this concept to the data in **Fig. 1E**, at 24 hpi, we found vDNA_min_ for TB40_Fb_ infecting epithelial cells to be 0.25 genomes/cell (vDNA_in,0_ = 0.3) and 0.84 genomes/cell (vDNA_in,0_ = 1) representing 26-42% reductions, and for TB40_Epi_ infecting epithelial cells to be 0.26 genomes/cell (vDNA_in,0_ = 0.6) and 1.2 genomes/cell (vDNA_in,0_ = 3) representing 55-58% reductions. The biological relevance of these differences remains unknown but could reflect differences in viral sources (e.g., stock infectivity, free vDNA, etc.) and entry mechanics.

To extend our analyses, we fit the empiric model of vDNA replication previously proposed (16) to each vDNA time course dataset (equations in **Fig. 2A**). We simplified the fitting process to allow for analysis of time courses of two vDNA_in,0_, rather than the previous three (16). Previously, nonlinear relationships were developed for each parameter (i.e., *vDNA_rep,max_*, *t_50_*, and *nH*) versus vDNA_in,0_, except for *k_d_* which was fixed (16). In this study, we estimated the value of *k_d_* (degradation rate), *t_50_*(time at which 50% of vDNA replication has occurred), and *nH* (empiric Hill coefficient) by fitting the equations in **Fig. 2A** to the mean time courses for each infection condition (i.e., virus source and target cell type) and vDNA_in,0_. Conversely, we maintained the functional form of vDNA*_rep,max_*as the saturable nature of this parameter was a key finding of (16). For fitting, the *V_max_* parameter (amount of vDNA at saturable vDNA_in,0_) in **Fig. 2A** was simultaneously fit to the time course for each vDNA_in,0_ in each condition along with *k_d_*, *t_50_*, and *nH* (**Fig. 2B**). The results of the simultaneous fitting to the time courses for both vDNA_in,0_ for each infection condition were then used in subsequent analysis. For consistency, this process was also repeated for the vDNA data from our previous studies (16). Fitting to mean data is shown in the lower panels of **Fig. 2C-E**, and for each individual replicate in the upper panels of **Fig. 2C-E**. Through this fitting process, we deduced that the *k_d_*, *V_max_*, and *nH* parameters (16) varied little between conditions (**Fig. 2B**). Although not significant, a notable difference exists between *k_d_* of TB40_Fb_ infection of fibroblasts versus epithelial cells (**Fig. 2B**). This suggests that the intrinsic characteristics of vDNA kinetics (i.e., degradation and replication) vary little between viral source-target cell combination. **Table 1** shows measured and simulated values for vDNA time courses. Overall agreement between measured and simulated values suggests accurate description of the data by this model.

**Figure 2.**
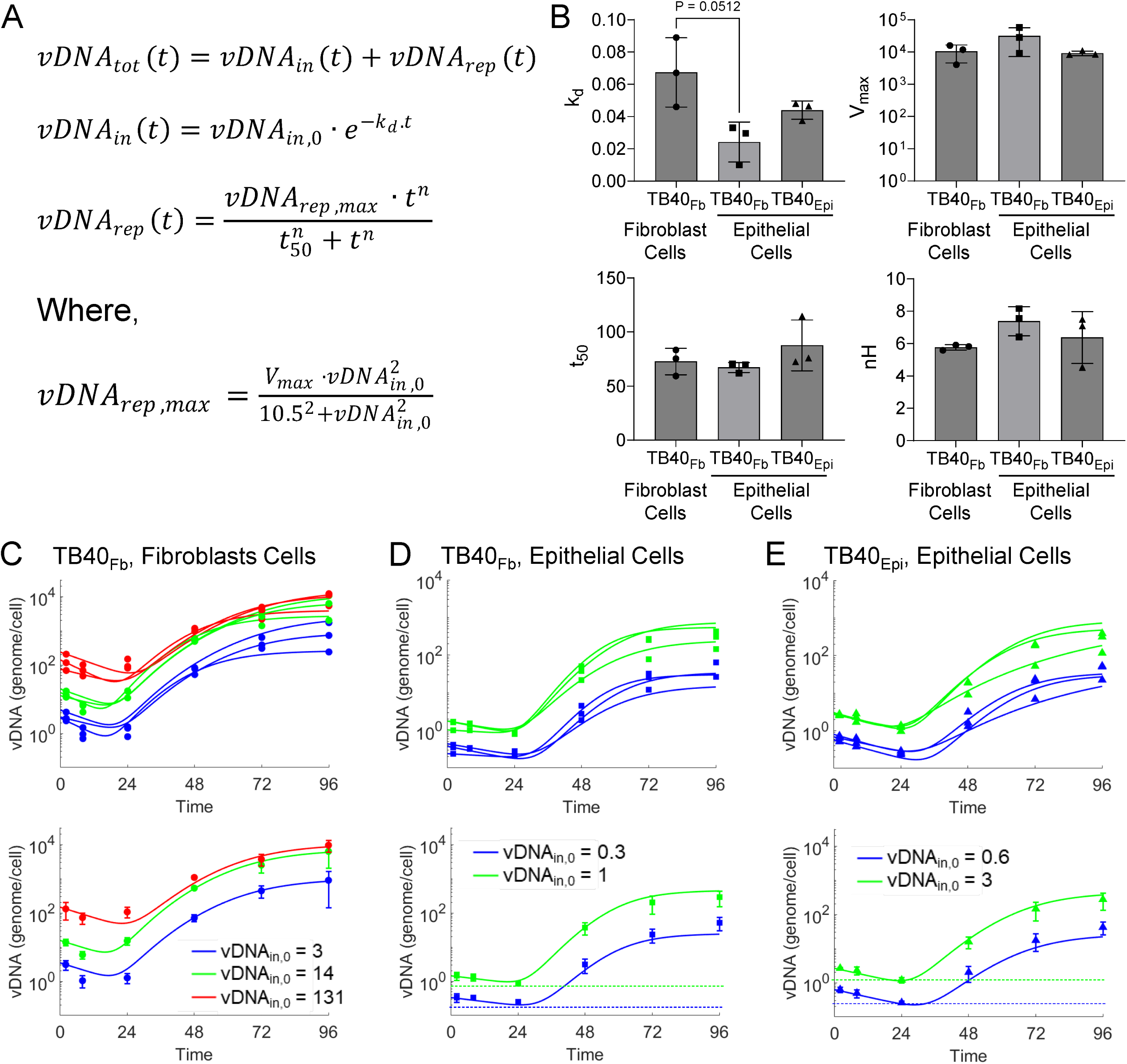
HCMV genome replication kinetics between different sources of virus during infection of epithelial cells. (**A)** Equations describing empirical vDNA model and estimation of vDNA_rep,max_ (16). **(B)** Parameters for vDNA replication estimated by fitting the model to individual biological replicate time courses. (*k_d_*, degradation rate; *V_max_*, amount of vDNA at saturable vDNA_in,0_; *t_50_*, time at which 50% of vDNA replication has occurred; and *nH*, empiric Hill coefficient; N = 3.) (**C**-**E**) Fit of vDNA model to individual time course data sets (upper panels) or mean time course data sets (lower panels) for **(C)** TB40_Fb_ on fibroblast (16), **(D)** TB40_Fb_ on epithelial cells, and **(E)** TB40_Epi_ on epithelial cells.

**Table 1.**
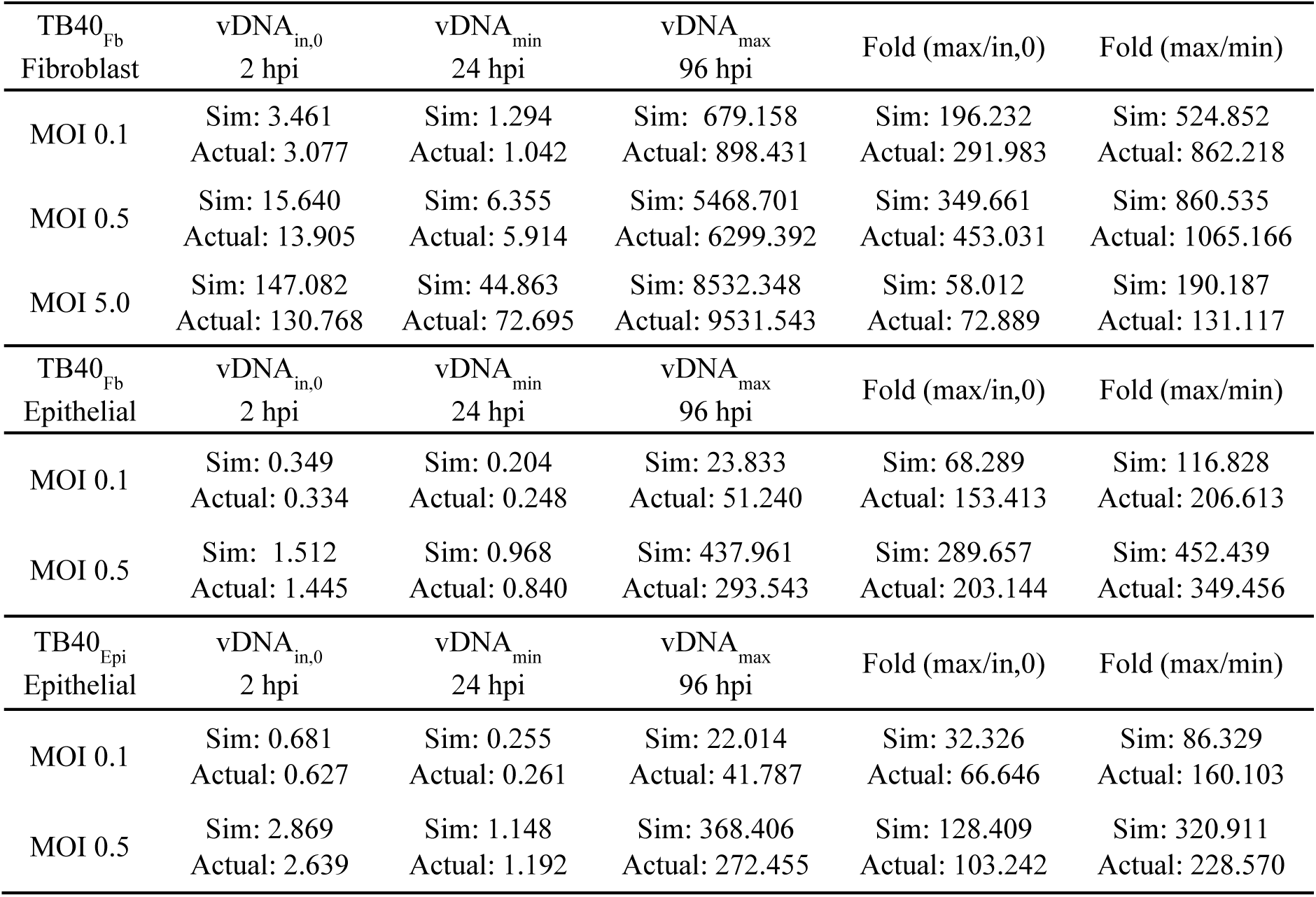
Experimental and simulated data for HCMV vDNA replication under multiple conditions.

We simulated vDNA replication kinetics for a wide range of vDNA_in,0_ (i.e., cell-associated vDNA at 2 hpi) varying between 10^-4^ genomes/cell to 10^4^ genomes/cell input. These simulations are shown in the 2D contours and 3D surfaces colorized with the fold change in vDNA at 96 hpi relative to vDNA_in,0_ (**Figs. 3A-C**). These *in silico* simulations suggest that there is little variability in vDNA kinetics between infection condition when varying vDNA_in,0_. This is in accordance with the findings in **Fig. 2B** with maximal fold change occurring at vDNA_in,0_ ∼10 genomes/cell which is consistent with our previous studies of TB40_Fb_ infection of fibroblasts (16). Also, high input at ∼1000 genomes/cell exhibit limited genome replication while very low input ∼1 genome/1000 cells show no replication likely with genome decay.

**Figure 3.**
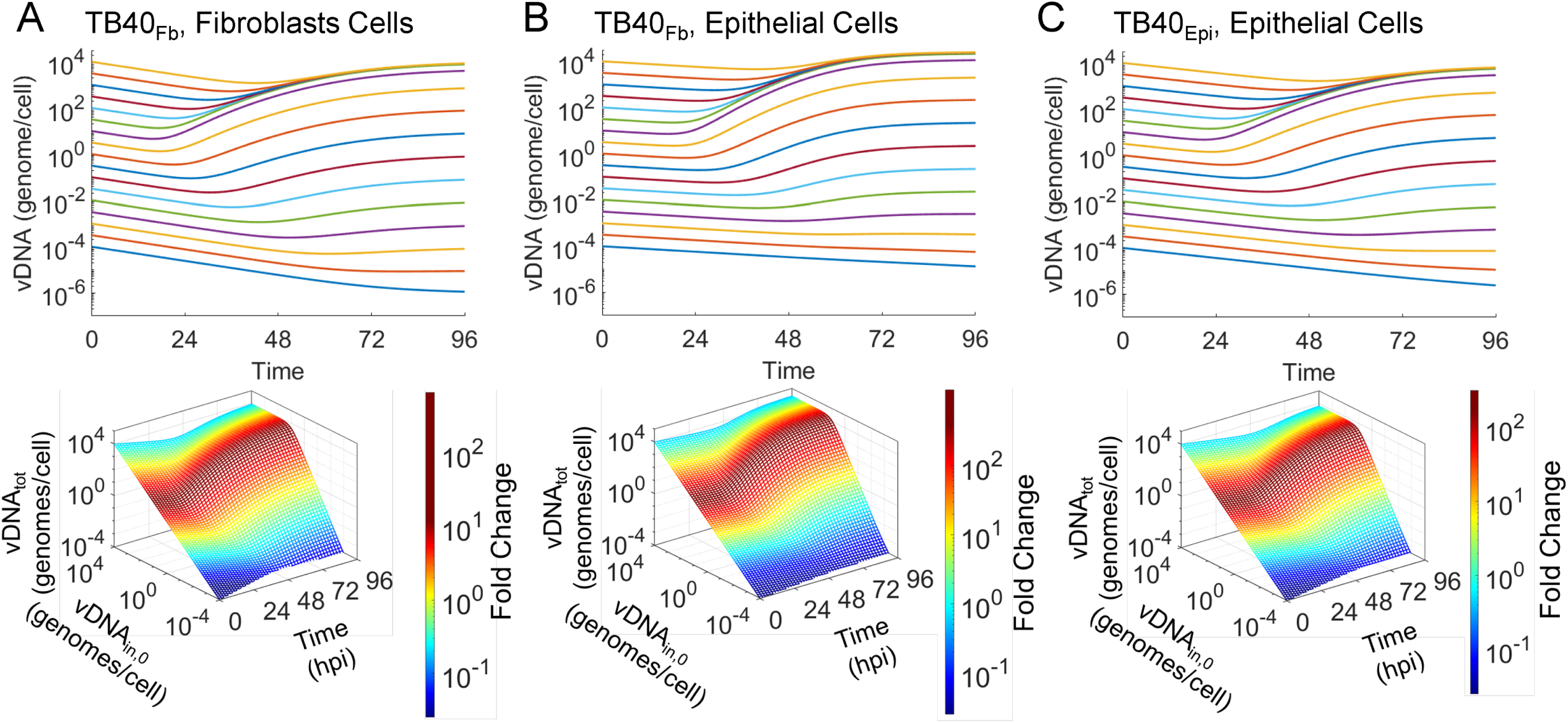
Simulations of vDNA replication show comparable kinetics between viral sources and target cell types. Simulations of vDNA replication kinetics at varying vDNA_in,0_ of 10^-4^ to 10^4^ vDNA/cell for **(A)** TB40_Fb_ on fibroblasts (16), **(B)** TB40_Fb_ on epithelial cells, and **(C)** TB40_Epi_ on epithelial cells. Three-dimensional plots (bottom panels) show relationships of vDNA_in,0_, vDNA_tot_, time, and fold change (vDNA (t = 96 hpi)/vDNA_in,0_) for source of virus. The optimal vDNA_in,0_ for each condition to achieve maximum fold change is indicated by a vertical line.

### Expression kinetics of Tp5 proteins show variability based on viral source during infection of epithelial cells

We continued to investigate the effect of viral source on the expression kinetics by evaluating two proteins in the late temporal class (Tp5), pUL44 and pp28 (10). We performed immunoblot analysis on whole cell lysates collected from the previously described experiments and probed for pUL44 and pp28 proteins. For comparison purposes only, the immunoblot for pUL44 and pp28 from our previously published studies are provided (**Fig. 4A, D**) (16). Importantly, to compare between blots we incorporated an internal standard and analyzed the data as previously described (16). Notably, we found that at the higher MOI, pUL44 levels trended higher in TB40_Fb_-infected epithelial cells compared to TB40_Epi_-infected cells (**Figs. 4B-C & G**). In contrast, pp28 levels trended higher in TB40_Epi_-infected epithelial cells compared to TB40_Fb_-infected epithelial cells (**Figs. 4E-F & H**). We next measured the amount of virus produced by ARPE19 cells infected with either TB40_Epi_ or TB40_Fb_ at two different MOIs (**Fig. 4I**). These data showed that TB40_Epi_ produced higher viral titers than TB40_Fb_ at both MOIs despite having similar viral genomes per cell at 96 hpi. Of note, prior to 72 hpi, there was an elevated amount of virus that progressively decreased suggesting the presence of residual input virus, and for subsequent computational modeling studies, 24 and 48 hpi data points were omitted as previously described (16). Given the similarity in vDNA kinetics between infection conditions, increased virus production from TB40_Epi_ relative to TB40_Fb_ in epithelial cells could be related to higher levels of pp28. These data suggest that the match of TB40_Epi_ viral infection with ARPE19 cells results in more efficient replication than unmatched TB40_Fb_ with ARPE19 cells, irrespective of vDNA replication kinetics.

**Figure 4.**
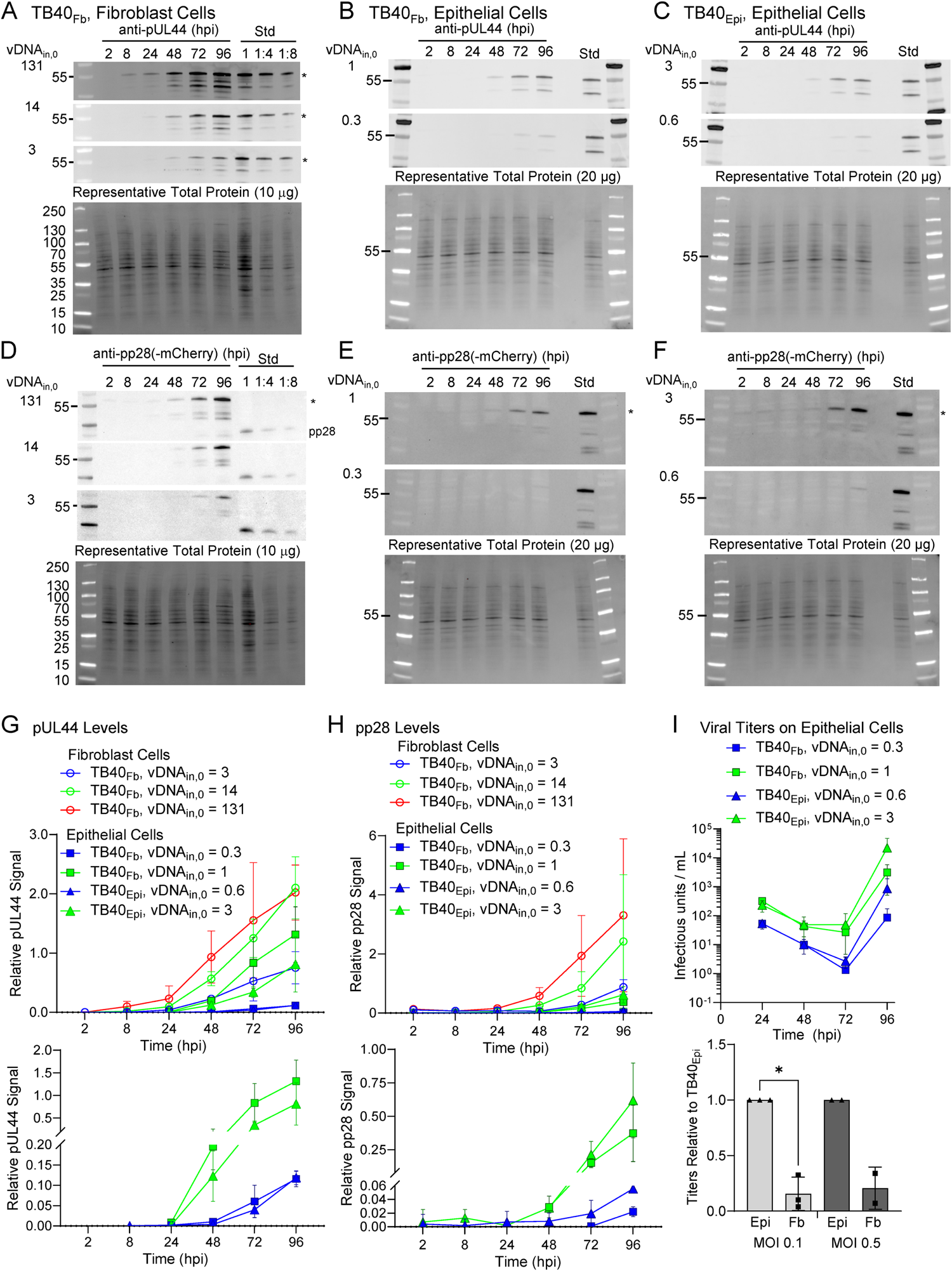
Expression kinetics of HCMV proteins pUL44 and pp28 during infection. Confluent ARPE-19 cells were infected with TB40_Fb_ or TB40_Epi_ at two input vDNA_in,0_ and done in parallel to studies quantifying vDNA. Whole cell lysates were isolated at the indicated time points and analyzed by immunoblot. Expression of pUL44 in **(A)** TB40_Fb_ on fibroblasts from previous studies (16), **(B)** TB40_Fb_ on epithelial cells, and **(C)** TB40_Epi_ on epithelial cells. Representative data are shown including total protein stains. Each immunoblot contains the same loading control standard (Std) allowing for comparisons between immunoblots. Similarly, expressions of pp28 are shown in **(D)** TB40_Fb_ on fibroblasts (16), **(E)** TB40_Fb_ on epithelial cells, and **(F)** TB40_Epi_ on epithelial cells. Signal volumes were obtained for each time point and normalized to total protein for a given sample set, and replicates were normalized to the standard run on each immunoblot. For epithelial cell studies, data were obtained from three biological replicate experiments for each virus source and vDNA_in,0_ with standard deviations are shown. The resulting quantifications are shown for **(G)** pUL44 and **(H)** pp28 with the lower panel expression in only epithelial cells. **(I)** Cell-free viral titers were measured on epithelial cells as IU/ml for TB40_Fb_ and TB40_Epi_ infections of epithelial at varying vDNA_in,0_. Data were obtained from three biological replicate experiments for MOI 0.1 and two for MOI 0.5. Due to variability between replicates, relative differences per replicate are shown and significance determined using a paired T-test.

### Modeling of late lytic replication reveals virus source-dependent kinetics of viral protein synthesis

Recently, computational modeling approaches have been used to reveal novel aspects of the HCMV lytic replication cycle (15, 16). This includes our postulated vDNA_in,0_-dependent model of late lytic replication that describes the major phenomena occurring during the final stages of HCMV lytic replication (schematic reproduced in **Fig. 5A** and equations reproduced in **Fig. 5B**) (16). In this model, total vDNA (vDNA_tot_) kinetics are described by an empirical input function (**Fig. 2**), which drives a mechanistic model of late Tp5 protein expression. The Tp5 proteins described by this model are subdivided into two categories: (1) nuclear-localized, capsid-associated (Tp5_1_) and (2) cytoplasmic, tegument-associated (Tp5_2_) proteins. After synthesis, the Tp5_1_ proteins combine with vDNA to form loaded capsids, which exit the nucleus and join with Tp5_2_ proteins to form intracytoplasmic viral particles. Finally, the intracytoplasmic particles leave the cell and form cell-free HCMV virions.

**Figure 5.**
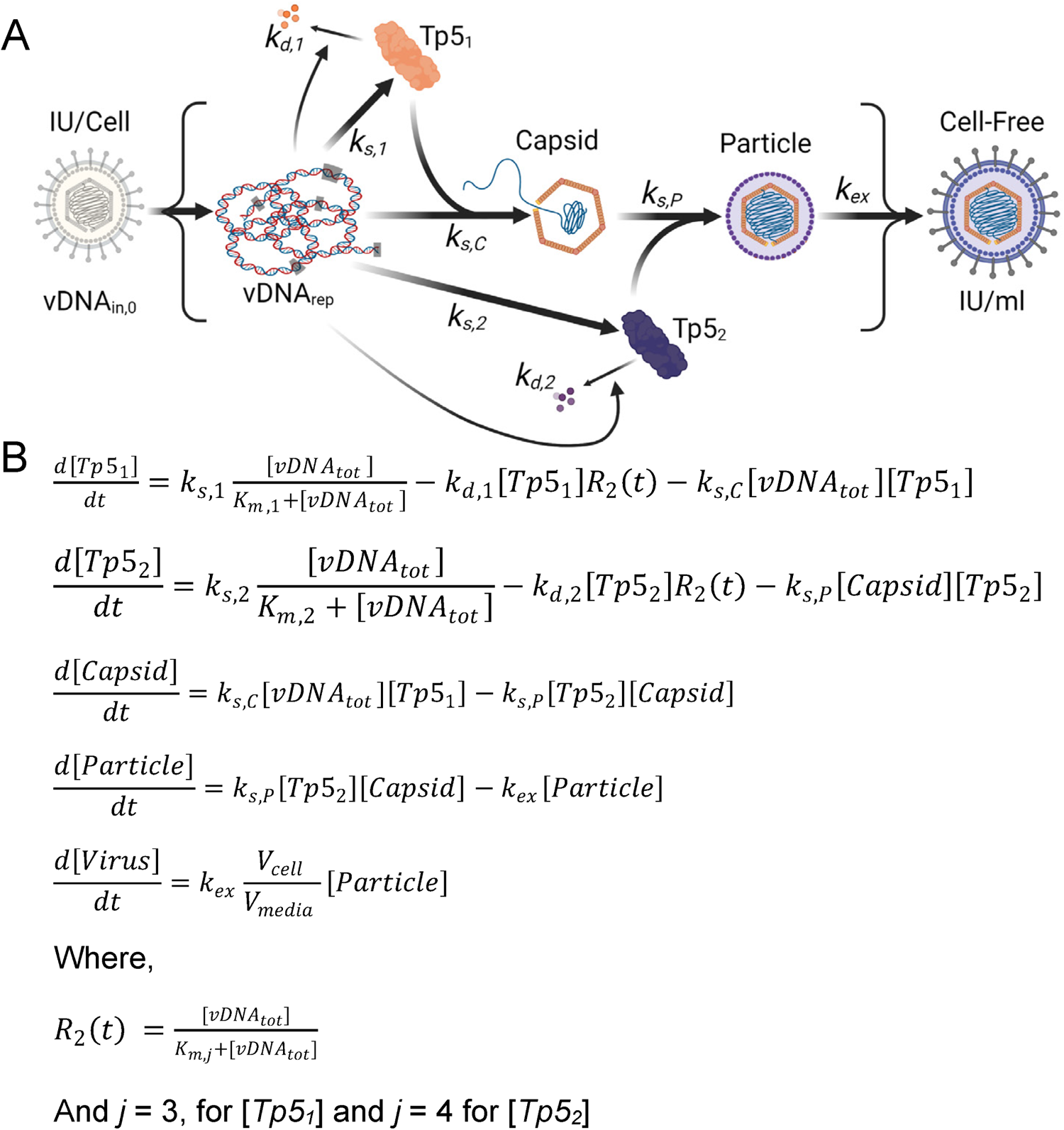
Postulated mechanistic model of HCMV late lytic replication cycle. (**A**) Model of late lytic HCMV replication by Monti et al. (16) developed and parametrized using experimental data from TB40_Fb_ infection of fibroblasts. HCMV infection defined by IU/cell (infectious units/cell) resulting in vDNA_in,0_ (input genomes measured at 2 hpi) and genome synthesis of vDNA_rep_ (replicated viral genomes) resulting in expression of nuclear protein Tp5_1_ (temporal profile 5) and cytoplasmic protein Tp5_2_, and subsequent assembly of capsid, intracellular particle and extracellular cell-free virus at IU/ml. Reactions are governed by various rates k_x_ (d, degradation; s, synthesis; s,C capsid synthesis; s,P particle synthesis; ex, virion egress). **(B)** Set of ordinary differential equations describing late lytic HCMV replication based on an empirical model of HCMV vDNA replication (V_cell_, cell volume; V_media_, media volume).

Using a pseudo-Monte Carlo parameter estimation approach, we fit the mechanistic model of late protein synthesis and viral egress (16) to the newly obtained protein expression data for Tp5_1_ and Tp5_2_ (**Fig. 4G, H**) and the viral titers (**Fig. 4I**). Protein expression data was renormalized as described in the methods section. Fittings of all viral protein and titer data are shown **Figure 6** including our previous studies for comparison (**Fig. 6A**) (16), TB40_Fb_ on epithelial cells (**Fig. 6B**), and TB40_Epi_ on epithelial cells (**Fig. 6C**). The output from the model provides viral titers in arbitrary units (A.U.). To convert titers to IU/mL, we multiplied the model outputs by the conversion factor 2×10^11^ IU/(A.U. mL). Experimental data points are overlayed with simulated line trajectories for each condition. **Table 2** compares individual experimental measurements and model simulations. Using our initial postulated vDNA_in,0_-dependent model, we observe some agreement between simulated and experimental data. This indicates that our basic framework (**Fig. 5**) incorporates some but not all mechanistic relationships necessary to precisely simulate virus production.

**Figure 6.**
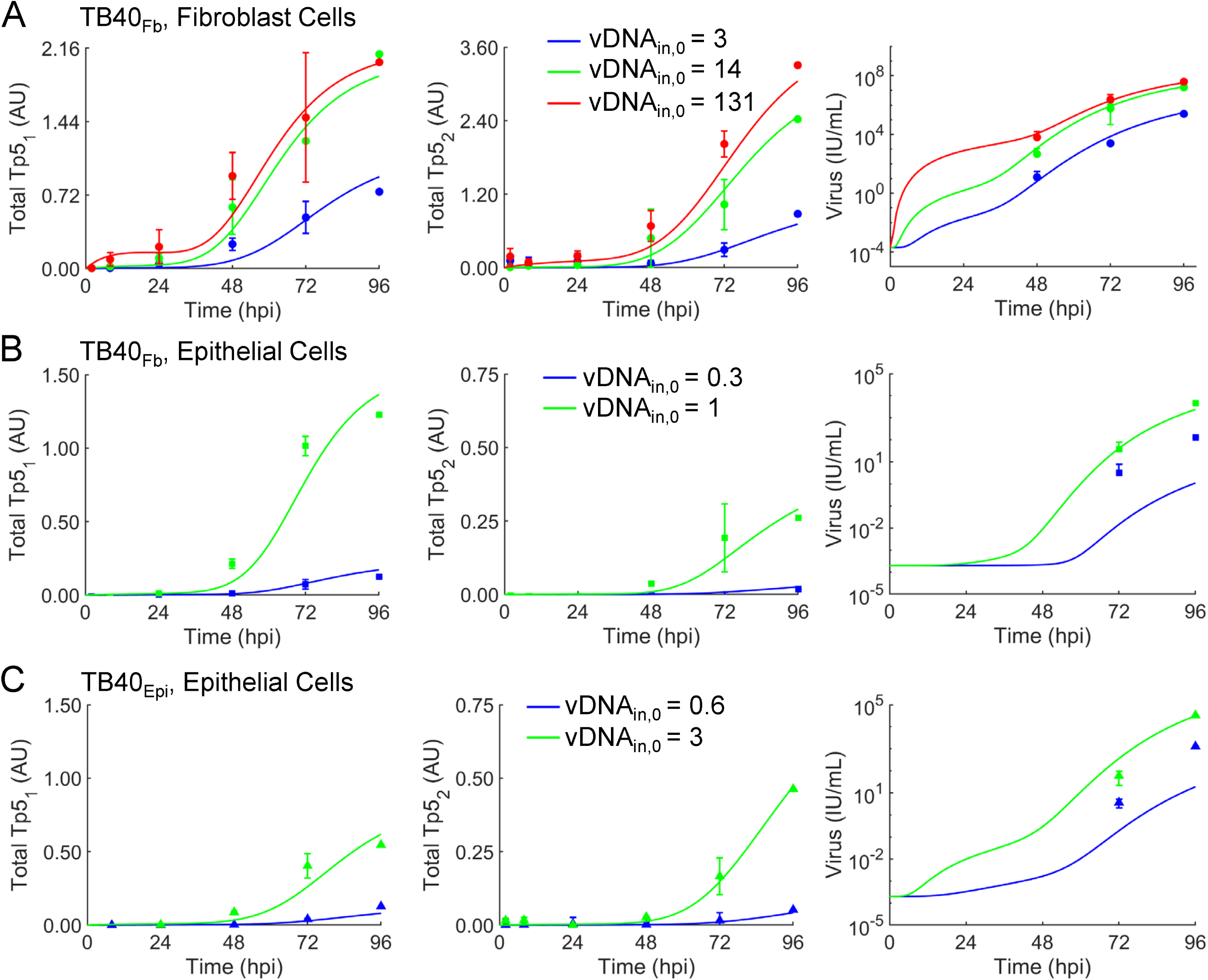
Altered kinetics of expression for late Tp5 proteins between TB40_Epi_ and TB40_Fb_ during infection of epithelial cells. Fit of model to immunoblot data for pUL44 (Tp5_1_) and pp28 (Tp5_2_) and viral titer data during infection using multiple vDNA_in,0_ of **(A)** TB40_Fb_ infection of fibroblast cells (16), **(B)** TB40_Fb_ on epithelial cells, and **(C)** TB40_Epi_ on epithelial cells. Time course datasets were first renormalized to minimize the error in the 96 hpi time points (see **Materials & Methods**). The ODEs for the model schematized were fit to each immunoblot dataset for pUL44 (Tp5_1_) and pp28 (Tp5_2_) and titering dataset (markers) using a 1000-iteration, pseudo-Monte Carlo parameter estimation protocol. Best-fit parameters were determined using the histograms in Figure 7. Simulations for each condition using these parameters are presented as solid line curves.

**Table 2.** Experimental and simulated data for HCMV Tp5 protein and vDNA levels under multiple conditions.

| TB40<sub>Fb</sub><br>Fibroblast | <sup>1</sup>vDNA<sub>in,0</sub> | <sup>1</sup>vDNA<sub>max</sub><br>96 hpi | Tp5<sub>1</sub> | Tp5<sub>2</sub> | Titer |
| --- | --- | --- | --- | --- | --- |
| MOI 0.1 | Sim: 3.461<br>Actual: 3.077 | Sim: 679.158<br>Actual: 898.431 | Sim: 0.899<br>Actual: 0.753 | Sim: 0.711<br>Actual: 0.878 | Sim: 3.121E5<br>Actual: 2.502E5 |
| MOI 0.5 | Sim: 15.640<br>Actual: 13.905 | Sim: 5468.701<br>Actual: 6299.392 | Sim: 1.887<br>Actual: 2.101 | Sim: 2.466<br>Actual: 2.425 | Sim: 1.768E7<br>Actual: 1.610E7 |
| MOI 5.0 | Sim: 147.082<br>Actual: 130.768 | Sim: 8532.348<br>Actual: 9531.543 | Sim: 2.012<br>Actual: 2.022 | Sim: 3.047<br>Actual: 3.307 | Sim: 3.326E7<br>Actual: 3.910E7 |
| TB40<sub>Fb</sub><br>Epithelial | vDNA<sub>in,0</sub> | vDNA<sub>max</sub><br>96 hpi | Tp5<sub>1</sub> | Tp5<sub>2</sub> | Titer |
| MOI 0.1 | Sim: 0.349<br>Actual: 0.334 | Sim: 23.833<br>Actual: 51.240 | Sim: 0.172<br>Actual: 0.124 | Sim: 0.026<br>Actual: 0.019 | Sim: 1.107E0<br>Actual: 1.290E2 |
| MOI 0.5 | Sim: 1.512<br>Actual: 1.445 | Sim: 437.961<br>Actual: 293.543 | Sim: 1.365<br>Actual: 1.228 | Sim: 0.291<br>Actual: 0.261 | Sim: 2.399E3<br>Actual: 4.570E3 |
| TB40<sub>Epi</sub><br>Epithelial | vDNA<sub>in,0</sub> | vDNA<sub>max</sub><br>96 hpi | Tp5<sub>1</sub> | Tp5<sub>2</sub> | Titer |
| MOI 0.1 | Sim: 0.681<br>Actual: 0.627 | Sim: 22.014<br>Actual: 41.787 | Sim: 0.080<br>Actual: 0.128 | Sim: 0.042<br>Actual: 0.052 | Sim: 1.899E1<br>Actual: 1.293E3 |
| MOI 0.5 | Sim: 2.869<br>Actual: 2.639 | Sim: 368.406<br>Actual: 272.455 | Sim: 0.618<br>Actual: 0.546 | Sim: 0.472<br>Actual: 0.463 | Sim: 3.112E4<br>Actual: 3.320E4 |

The raw output of the pseudo-Monte Carlo parameter estimation approach is a set of histograms showing the frequency of estimation for the value of each estimated parameter after 1,000 iterations (16). These histograms are shown in (**Fig. 7A**) for each condition. The most frequently estimated parameter was selected from these histograms to generate the curves in **Figure 6**. We compared the most frequently estimated parameters (**Fig. 7B**) for each fitting and found that the synthesis parameter (k_s,1_) for Tp5_1_ (pUL44) was higher for TB40_Fb_ infecting epithelial cells than TB40_Epi_ despite a lower vDNA_in,0_ (**Table 1**). The opposite was true for the synthesis parameter (k_s,2_) for Tp5_2_ (pp28) which was higher for TB40_Epi_ infecting epithelial cells than TB40_Fb_ and more closely followed the trends in vDNA_in,0_ and titers (**Tables 1** and **2**). It is important to note that the intrinsic Michaelis-Menten parameters for synthesis of Tp5_1_ (K_m,1_) and Tp5_2_ (K_m,2_) were fixed to values estimated from our previously fitting of the data for TB40_Fb_ infecting fibroblast cells (16), since a rise in synthesis parameter could be counteracted by a concomitant increase in the Michaelis-Menten parameter. Given the limited data for TB40_Epi_ and TB40_Fb_ infecting epithelial cells, we opted to fix these parameters to the values estimated from fitting TB40_Fb_ infecting fibroblast cells as these parameters had been estimated in previous studies and our results could be qualitatively compared to those previously established (16). Finally, k_s,P_ (viral particle membrane coating and egress from infected cell) and k_ex_ (intracellular particle release into extracellular media) showed higher rates during TB40_Epi_ replication in ARPE19 cells (**Fig. 7B**) which are consistent with experimental titer data (**Fig. 4I**). It is important to note that titers from using TB40_Fib_ on fibroblasts resulted in higher titers (16) at comparable vDNA_in,0_ which is reflected in a high k_ex_ compared to infection of epithelial cells (**Fig. 7B**). Finally, we simulated values of all predicted (free Tp5_1_, free Tp5_2_, Capsid, Particle, and Virus in A.U.) and derived measurable state variables (total Tp5_1_, total Tp5_2_, and virus in IU/mL) for vDNA_in,0_ ranging from 10^-4^ to 10^4^ genomes/cell (**Fig. 8**). For comparison, simulations from (16) are provided (**Fig. 8A**) along with TB40_Fb_ on epithelial cells (**Fig. 8B**) and TB40_Epi_ on epithelial cells (**Fig. 8C**). These simulations show consistently elevated Tp5_1_ in TB40_Fb_ infection over TB40_Epi_, elevated Tp5_2_ in TB40_Epi_ over TB40_Fb_ along with increased viral titers at comparable vDNA_in,0_. These data also show similar trends as shown previously including saturation kinetics with large increase in Tp5_1_ and Tp5_2_ production early in infection at high MOI or vDNA_in,0_ (16).

**Figure 7.**
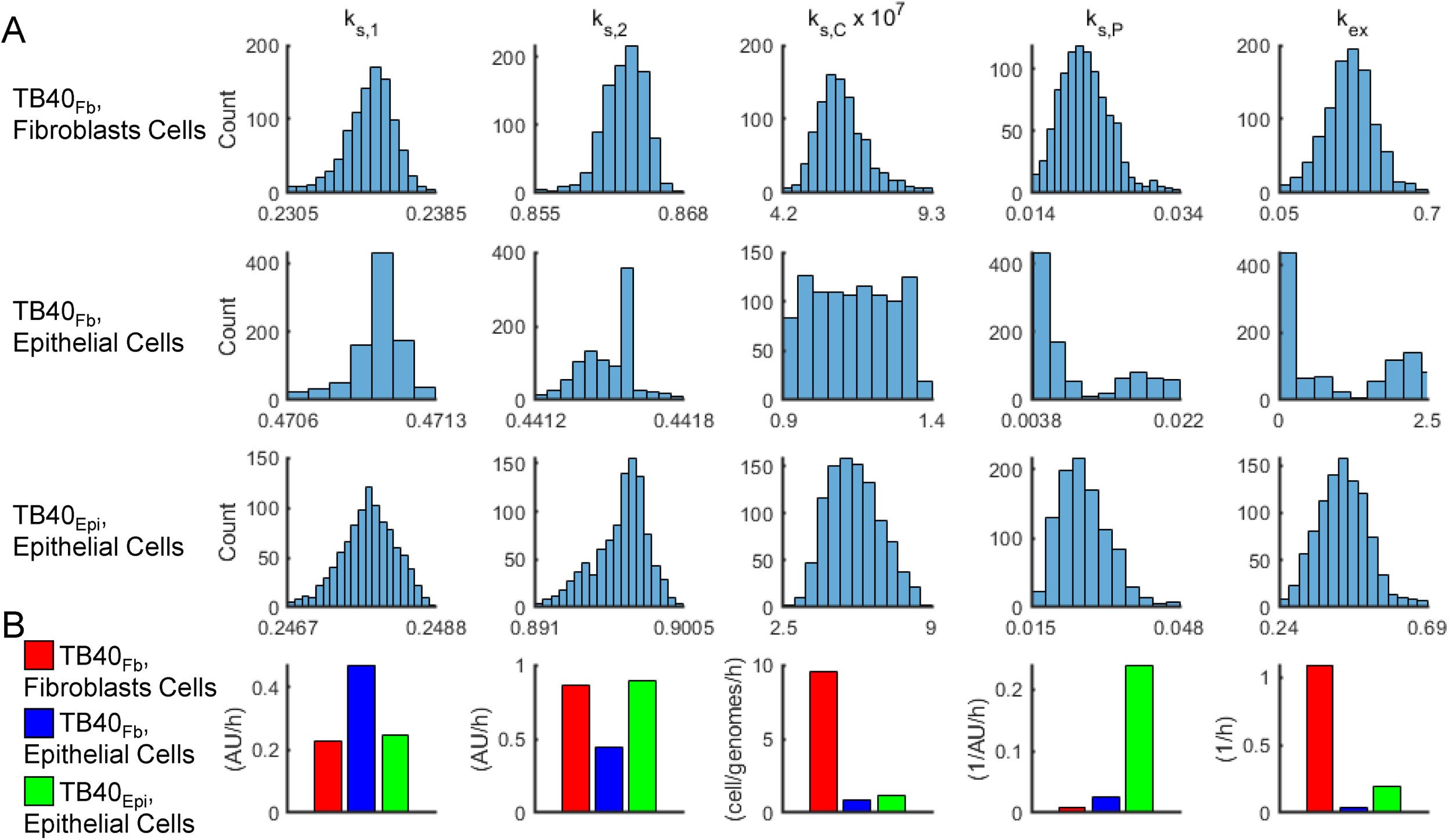
Parameter estimates using the pseudo-Monte Carlo fitting strategy for each infection condition. **(A)** Histograms of parameter estimates for 1000 iterations of the pseudo-Monte Carlo parameter estimation protocol. Estimated parameter values were collected during each iteration of the parameter estimation protocol. Each of the 1000 estimates for all parameters was binned and a count of the number of estimates in each bin was plotted using the MATLAB histogram function. **(B)** Comparison of model parameter estimates using the mean of most frequently estimated value ±20%.

**Figure 8.**
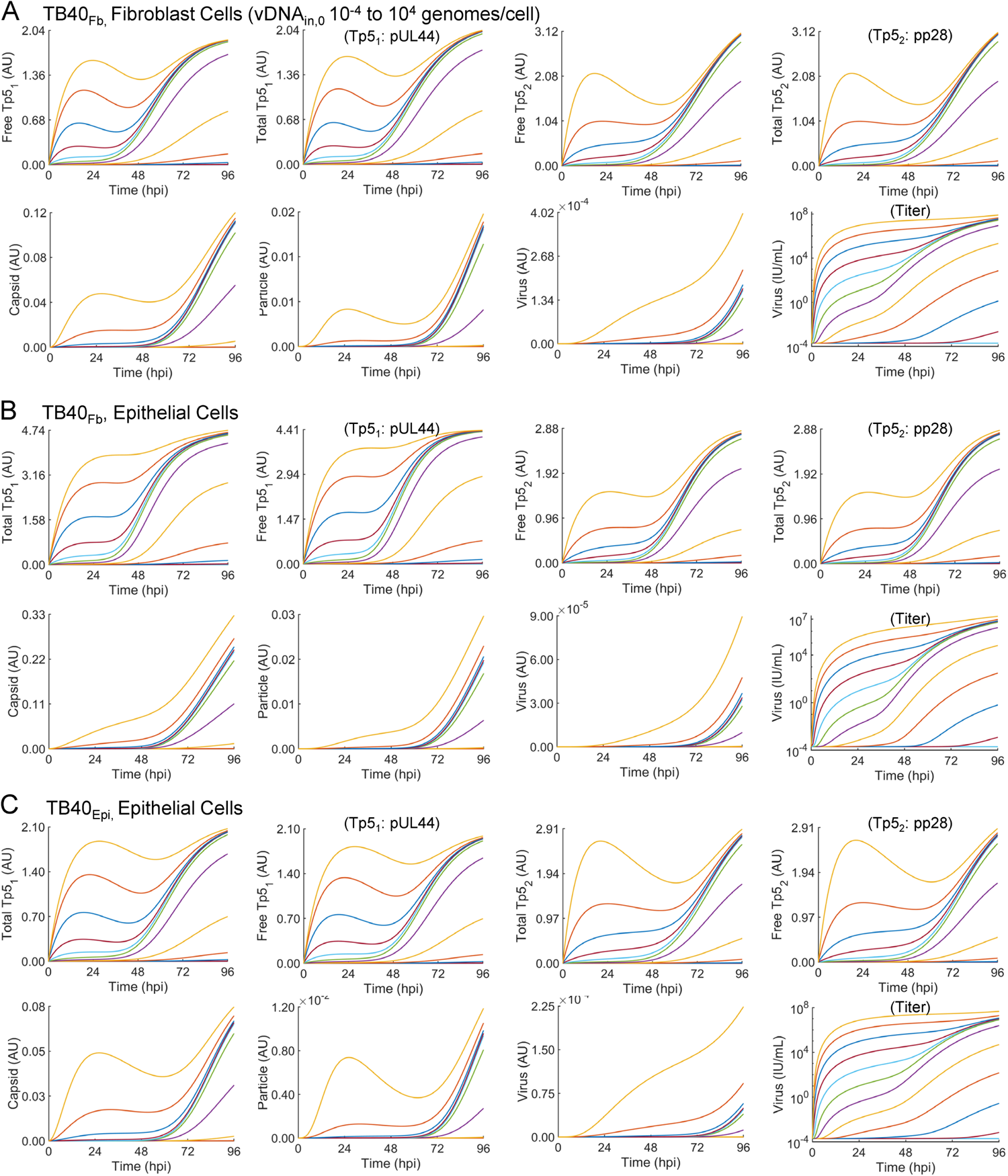
Simulations of late lytic replication for each infection condition at many vDNA_in,0_. Simulations of predicted (i.e., not fit) state variables free Tp5_1_, free Tp5_2_, Capsid, Particle, and output viral titers in units of AU and experimentally derived (i.e., fit) state variables total Tp5_1_, total Tp5_2_, and viral titers in units of IU/mL for vDNA_in,0_ ranging from 0.0001 to 10,000 vDNA/cell for **(A)** TB40_Fb_ infecting fibroblast, (**B**) TB40_Fb_ infecting epithelial cells, and (**C**) TB40_Epi_ infecting epithelial cells. From the ODE-based model, output viral titers are given in arbitrary units (AU). A conversion factor of 2 x 10^11^ IU/(AU mL) was applied to express output titer simulations in IU/ml to each plot yielding titer simulations in IU/ml.

### Epithelial viral source leads to syncytia formation and elevated pp28 levels when infecting epithelial cells

To further validate our experimental and computational findings, we infected APRE19 cells with TB40_Epi_ or TB40_Fb_ at MOI 0.1 and 0.5 and measured fluorescence intensity from tagged proteins IE2 (T2A-eGFP, green channel) and pp28 (pp28-mCherry, red channel) at a higher temporal resolution than achievable through immunoblot analysis using live cell imaging. Still images of TB40_Fb_-infected epithelial cells (**Fig. 9A**) show the prototypical pp28-mCherry cytoplasmic assembly compartment with diffuse free GFP signal by 42 hpi and increasing by 96 hpi. In TB40_Epi_-infected epithelial cells, cells exhibiting a similar pattern of mCherry with GFP are observed (**Fig. 9B**). However, and to our surprise, a second pattern of fluorescence was evident, consisting of larger pp28-mCherry signal intensities and areas with little GFP (**Fig. 9B**). A widefield view is also provided depicting these differences (**Fig. 9A, B**). To measure these differences, total fluorescence intensity for each channel was determined and plotted over time. This revealed higher GFP levels in TB40_Fb_-infected cells versus TB40_Epi_ over time, yet higher pp28-mCherry levels in TB40_Epi_-infected epithelial cells versus TB40_Fb_ (**Fig. 9C**). These observations are consistent with immunoblot data (**Fig. 4**) used in model parameterization and evident in simulated infections over a range of inputs (**Fig. 8**).

**Figure 9.**
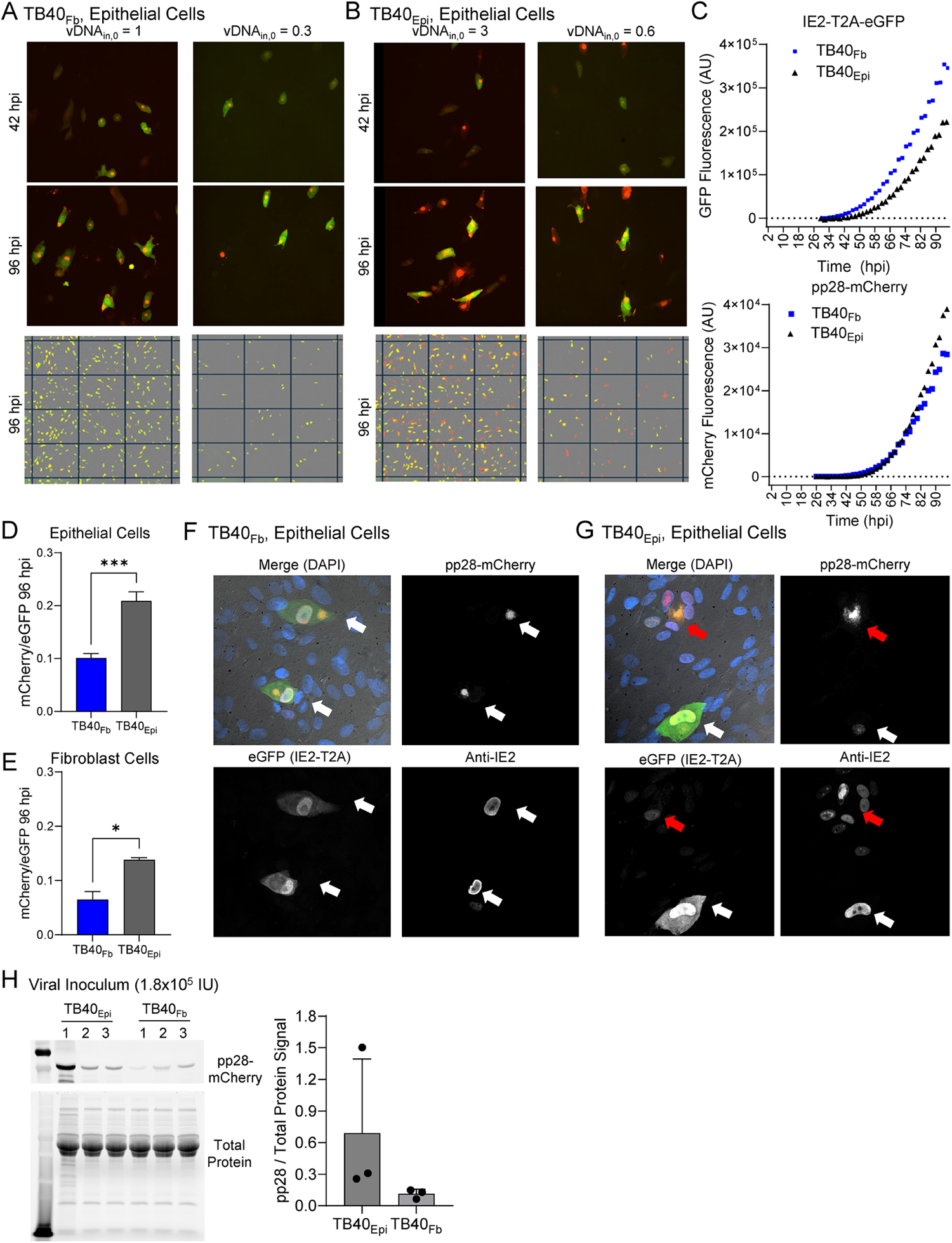
Infection of epithelial cells by TB40_Epi_ virus exhibits altered replication compartments at late times. Confluent ARPE-19 cells were infected at the indicated vDNA_in,0_ using **(A)** TB40_Fb_ or **(B)** TB40_Epi_ viruses. Infections were imaged every 2 hr for eGFP (IE2-T2A-eGFP) and pp28-mCherry until 96 hpi. Representative images are shown at 42 hpi and 96 hpi from three biological replicate experiments. A larger field of view is provided for each condition at 96 hpi. **(C)** Total integrated intensity for green channel (GCU x mm^2^/image) and red channel (RCU x mm^2^/image) over time from MOI of 0.5. The average background fluorescence prior to 24 hpi has been removed. Signal from TB40_Fb_ shown in blue squares and TB40_Epi_ in black triangles. **(D)** Ratio of mCherry to eGFP signals at 96 hpi following infection of epithelial cells using three independent viral stocks. **(E)** Ratio at 96 hpi following infection of fibroblasts by TB40_Fb_ or TB40_Epi_. **(F)** Representative confocal microscopy images at 96 hpi by TB40_Fb_ showing IE2 expression (eGFP or anti-IE2 antibody), pp28-mCherry, and merged images including Hoechst, and **(G)** by TB40_Epi_. Single cell replication compartments indicated by white arrows and altered replication compartments likely syncytia indicated by red arrows. (**H**) Comparison of pp28 levels from viral innocula. Volumes for 1.8×10^5^ IU of three different stocks of TB40_Epi_ or TB40_Fb_ were subjected to immunoblot analysis using an anti-pp28 antibody. Quantification of band intensity normalized to total protein are shown for TB40_Epi_ and TB40_Fb._ virus stocks.

We next asked whether these differences were viral source- or target cell type-dependent by measuring the ratio of mCherry to eGFP signal at the 96 hpi time point. As expected, this ratio was significantly higher upon TB40_Epi_ infection of epithelial cells (**Fig. 9D**). We used different sources of virus to infect MRC-5 fibroblasts during live-cell imaging which also showed a trend of a higher ratio in TB40_Epi_ versus TB40_Fb_ (**Fig. 9E**). These data indicate that the expression differences are more dependent on the virus source. We evaluated the infected cells using confocal microcopy which again shows the prototypic staining in the TB40_Fb_-infected epithelial cells of a cytoplasmic pp28-mCherry assembly compartment (**Fig. 9F**). Since eGFP is cleaved from IE2, we also co-stained using an antibody for IE2 which localized to the nucleus of infected cells. We repeated this approach following TB40_Epi_ infection of epithelial cells. This revealed two types of staining patterns, the prototypical pattern and a pattern consisting of a larger pp28-mCherry replication compartment with low eGFP signal (**Fig. 9G**). Co-staining with anti-IE2 shows lower levels of IE2 in smaller nuclei surrounding the assembly compartment with this organization suggestive of syncytia formation. Finally, to test whether there might be general differences in protein composition between TB40_Fb_ and TB40_Epi_ viruses, we completed an immunoblot analysis on viral inoculum from three different viral stocks for each source using 1.8×10^5^ IU per lane and anti-pp28 antibody (**Fig. 9H**). We quantified the relative amount of pp28 signal normalized to total protein which showed a trend toward higher pp28 levels in TB40_Epi_ compared to TB40_Fb_ virus. These data indicate variability in virion composition occurs when produced from different cell types.

Finally, using the updated simulations of the late lytic replication cycle, we evaluated the impact of varying vDNA_tot_ at 96 hpi on viral titers at 96 hpi. These results suggest that infection of epithelial cells by TB40_Epi_-sourced virus requires less vDNA_tot_ at 96 hpi compared to TB40_Fb_-sourced virus to produce similar amounts of infectious virus (**Fig. 10A**). This is supported in simulating varying vDNA_in,0_ from 10^-4^ to 10^4^ genomes/cell where TB40_Epi_ infection is predicted to produce higher viral titers per genome (**Fig. 10B**) and per Tp5_2_ (pp28) expression levels (**Fig. 10C**). Overall, these data indicate that the viral source is influencing the composition of the particles and subsequently influencing replication efficiency and perhaps the type of replication compartments (i.e., prototypic single cell verse syncytia) during HCMV infection.

**Figure 10.**
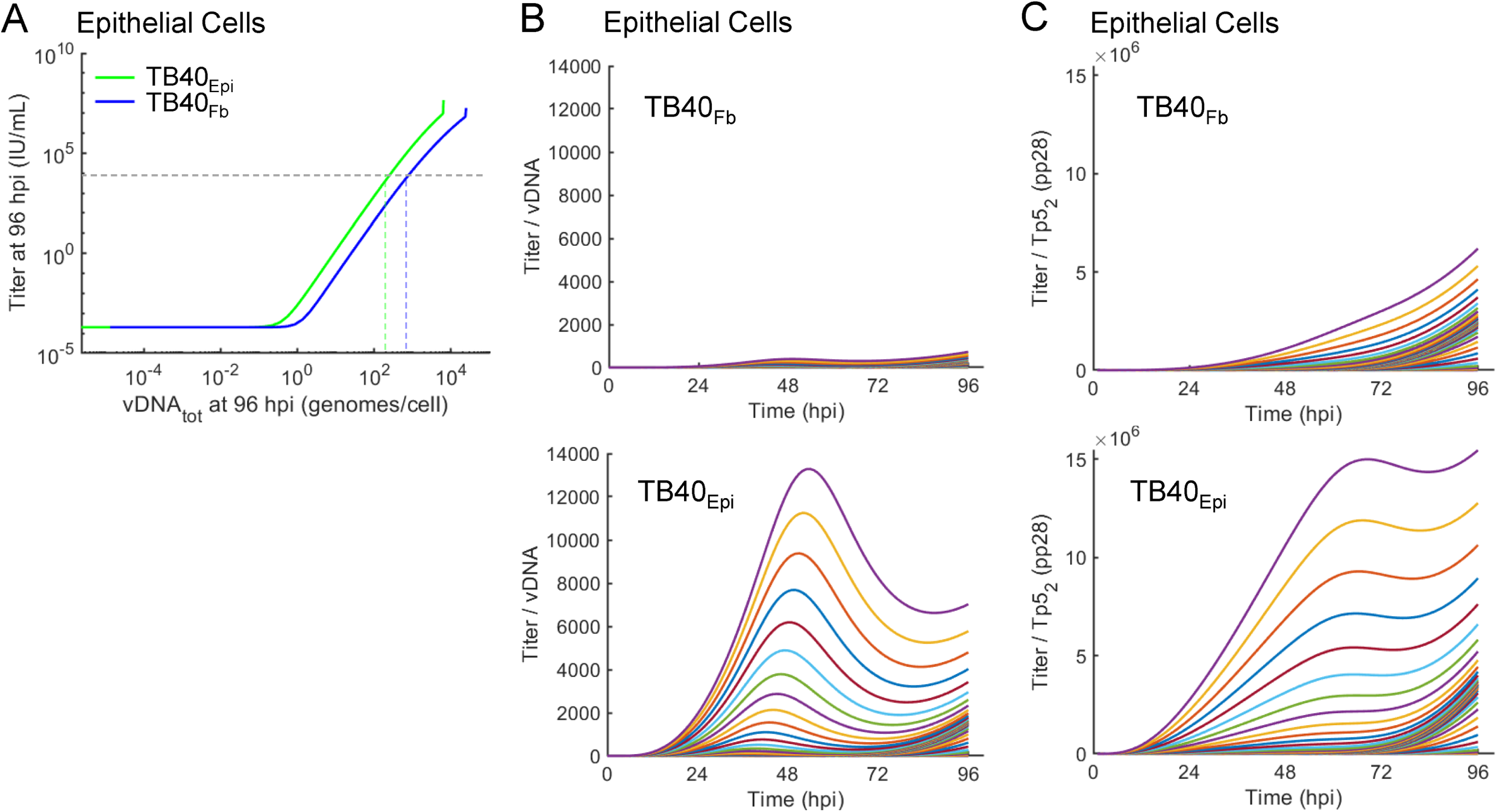
Comparison of simulated TB40_Epi_ and TB40_Fb_ replication efficiencies during infection of epithelial cells. **(A)** Comparison of simulated changes in viral titers versus vDNA_tot_ at 96 hpi for varying vDNA_in,0_ for TB40_Fb_ and TB40_Epi_. Dotted lines are added to visualize the intersection of 1×10^4^ IU/ml with vDNA_tot_ for each virus. (**B**) Simulations of changing ratio of titer/vDNA over time at multiple vDNA_in,0_ from 0.0001 to 10,000 IU/cell for TB40_Fb_ and TB40_Epi_. **(C)** Simulations of changing ratio of titer/Tp5_2_ (pp28) over time at multiple vDNA_in,0_ from 0.0001 to 10,000 IU/cell for TB40_Fb_ and TB40_Epi_.

## DISCUSSION

In this manuscript, we applied the combined experimental and computational modeling framework devised in our previous studies (16) to investigate the effects of virus source and target cell type on HCMV late lytic replication kinetics. In general, we found that the developed computational model describes the kinetics of viral DNA (vDNA), two late (Tp5) proteins, and viral titers determined experimentally, with fine tuning of model parameters required to obtain an acceptable fit of the model to the data. Using data-driven computational modeling along with additional experimental validation studies, we found that: (1) the parameters describing vDNA synthesis were similar between TB40_Fb_ (fibroblast-derived virus) infecting epithelial cells and TB40_Epi_ (epithelial-derived virus) infecting epithelial cells; (2) TB40_Epi_ infection of epithelial cells led to a larger amount of pp28 expression and higher titers in comparison to TB40_Fb_ infection of epithelial cells; and (3) the source of the virus impacts the replication efficiency (i.e., titer per genome and Tp5) likely involving altered virion composition and type of replication compartments (i.e., single-cell versus syncytia).

HCMV lytic replication is a complex and protracted process involving many viral and host proteins that combine in a specific and regulated network to produce progeny virions. The source of the viral particle and the type of the cell infected are two of many factors that can influence lytic replication. In this manuscript, we applied our previously developed computational model using one cell type and one viral source (16) to a new cell type, epithelial cells, and two different viral sources, TB40_Fb_ and TB40_Epi_. Specifically, we first fit the empirical model of vDNA replication (16) to time course qPCR data using absolute standards from epithelial cells infected with TB40_Epi_ or TB40_Fb_ (**Figs. 1-4**). By fitting the empiric vDNA replication model to individual time courses, we demonstrated that kinetic parameters were similar between the infection conditions. Overall, this suggests that the kinetics of vDNA replication are intrinsic to the HCMV virus regardless of viral source (epithelial cells or fibroblasts). However, we did observe variation between vDNA_in,0_ (vDNA/cell at 2 hpi) and vDNA_min_ (lowest level) as well as variation between MOIs and vDNA_in,0_. It is conceivable that these differences are partly due to technical variation in viral stock preparations and titering even when using the same methodology, altered stock infectivity (i.e., IU/genomes or IU/particle), and presence of cell-free viral DNA associating with cells. Alternatively, the rate from vDNA_in,0_ to vDNA_min_ (noted here as k_d_) may reflect altered mechanisms of entry with micropinocytosis, endocytosis, and cell surface fusion being previously demonstrated mechanisms (25–29). Future studies will focus on refining this rate as it has significant implications on understanding HCMV tropism and persistence.

We obtained multiple experimental datasets over time and two MOIs using three biological replicates to parameterize the model. We obtained immunoblot data describing the kinetics of two HCMV proteins, nuclear pUL44 and cytoplasmic pp28 (**Fig. 4**) which are defined by Weekes et al. (10) as belonging to Tp5 classification. We fit these data to our mechanistic model using a pseudo-Monte Carlo fitting strategy that had a relatively narrow range of estimated parameters and that conformed to the known and hypothesized constraints of the system while also qualitatively describing the data (**Fig. 5**). Like the empirical model for vDNA replication, the mechanistic model fits to the general kinetic trends for Tp5 with fine tuning of model parameters. We also measured the output viral titers from TB40_Epi_- or TB40_Fb_-infected epithelial cells (**Fig. 4I**) and simultaneously fit these data to the output of the model (**Figs. 5**, **6**). Using this approach, we uncovered a larger synthesis constant for pp28 expression during TB40_Epi_ infection of epithelial cells compared to TB40_Fb_. This elevated level of pp28 is experimentally evident in both immunoblot analyses (**Fig. 4**) and analyses of live-cell imaging data (**Fig. 9**). We found a higher pp28-mCherry signal in epithelial cells infected with TB40_Epi_ compared to TB40_Fb_. Interestingly, we also found that this trend was similar in MRC5 fibroblasts which implicates differences being dependent on the viral source. Purified HCMV virions contain up to 82 different viral proteins and at varying levels (30–32), and passage of BAC-derived virus in epithelial cells has been shown to alter virion protein levels, including altered ratios of trimeric gH/gL/gO versus pentameric gH/gL/pUL128-131 entry receptors (33). Therefore, we tested whether differences might be evident in our viral stocks by measuring tegument protein pp28 levels. These studies revealed higher levels of pp28-mCherry in TB40_Epi_ over TB40_Fb_ in sorbitol pelleted stocks at equal IU relative to total protein and from multiple stock preparations (**Fig. 9H**). Overall, our data support the idea that altered protein composition in virions likely exists between TB40_Epi_ and TB40_Fb_ viral stocks, contributing to differences in HCMV replication kinetics.

While our study is broadly based on the integrated experimental-computational framework defined in (16), there are still several limitations that must be addressed. First, while our integrated experimental and computational techniques have a solid foundation in the literature (e.g., (15, 16, 34, 35)), predictions made by the computational models require experimental validation. These models are hypothesis-generating, and our future studies will experimentally test these model predictions. A second limitation is the limited number of viral proteins being measured and the semi-quantitative, low-resolution nature of the data obtained from the immunoblot analysis. The increase in Tp5_2_ protein pp28, for example, could be investigated further by quantitating total pp28 using multiple-reaction monitoring mass spectrometry (MRM/MS) including absolute peptide standards as done for qPCR (36–38). Additionally, as the model proposed in (16) assumes that pp28 is a prototypical member of a broader class of Tp5_2_ proteins and thus additional proteins involved in tegument formation (e.g., pp65, pp71) could also be investigated using MRM/MS (39) to determine if the increase in pp28 is part of a more global increase in the Tp5_2_ family of proteins. Inclusion of additional viral proteins using higher resolution data will improve the predictive power of this model. Following further refinement, it will be important to investigate numerous cell types, and it will be particularly interesting to determine the kinetics of vDNA, Tp5 proteins, and viral titers in a model of HCMV latency and reactivation. On a similar note, it would also be important to develop a standardized protocol for infection using vDNA_in,0_ as opposed to MOI. As demonstrated in **Fig. 1**, infection at the same MOI can lead to differing vDNA_in,0_, which can make a large difference in lytic replication kinetics, as exhibited in the 3D plots in **Fig. 4**. Thus, moving forward, it is reasonable to suggest that investigators seeking to utilize computational modeling in their studies develop a consistent methodology to infect based on vDNA_in,0_ rather than MOI. While these limitations do present opportunities for further study, we do not think that they appreciably impact the broader findings of this manuscript.

Following infections using live-cell imaging, we observed infected cells showing prototypic single cell infections with deformed IE2-positive nuclei juxtaposed to a cytoplasmic assembly compartment and this occurred in both TB40_Epi_ and TB40_Fb_ infections (**Fig. 9F,G**). However, in TB40_Epi_ infections, we also observed a different organization consisting of larger areas of pp28-mCherry with little GFP signal. Upon analysis by confocal microscopy, we detected multiple IE2-positive nuclei with expression occurring at low levels surrounding a pp28-positive assembly compartment which is suggestive of syncytia. We recently demonstrated robust syncytia formation following TB40_Epi_ infection of neural progenitor cells and terminally-differentiated neurons (40, 41). Vo et al. (42) demonstrated that the original TB40/E virus stock adapts to infection of ARPE19 cells within two passages which includes development of multinucleated syncytia. The authors speculate that passage in epithelial cells results in improvements in entry efficiency. In our studies, we passaged TB40_Fb_ derived from TB40-BAC4 three times in epithelial cells in preparing TB40_Epi_. We detected similar kinetics of vDNA replication between viral sources yet measured subsequent differences in pp28 expression kinetics and organization within infected cells. It is unclear if these differences are related to increased entry efficiency; however, our data implicate differences in replication between the two sources of virus. In **Fig. 10**, we compare simulations of the relative amounts of output virions to vDNA and pp28 over time for TB40_Epi_ and TB40_Fb_ infecting epithelial cells between 10^-4^ to 10^4^ IU/cell. These plots suggest that for any vDNA_in,0_, the ratio of titer to vDNA or to pp28 is larger in TB40_Epi_ versus TB40_Fb_ when infecting epithelial cells. Our recent studies suggest that fibroblast cells infected by TB40_Fb_ have a maximal capacity to produce virus (16). With this concept in mind, we suggest that syncytia formation bypasses this single cell capacity allowing more effective usage of viral components to form new virions, hence the higher viral titers for TB40_Epi_ versus TB40_Fb_ at similar levels of vDNA. Finally, it is important to note the resulting titers from epithelial cells in our current studies remain lower than titers from infection of fibroblasts with TB40_Fb_ using comparable input vDNA (16).

In summary, we demonstrate that the virus source impacts the replication efficiency for the specific virus-cell combinations tested and likely involves altered input virion composition and resulting type of replication compartments. Inclusion of additional viral proteins using higher resolution data in future studies will improve the predictive power of this model as we move toward a complete mechanistic model of HCMV lytic replication.

## MATERIALS AND METHODS

### Cells, Viruses, and Biological Reagents

Dual fluorescently tagged TB40/E HCMV expressing IE2-2A-eGFP and UL99-mCherry was generously provided by Dr. Eain Murphy. Viral stocks were propagated as a P1 stock on MRC-5 fibroblasts (ATCC) or ARPE19 epithelial cells (ATCC) and concentrated by collecting culture medium and pelleting through a sorbitol cushion containing 20% D-sorbitol, 50 mM Tris-HCl pH 7.2, 1 mM MgCl_2_. The viral media overlayed on the sorbitol cushion was applied to a Sorvall WX-90 ultracentrifuge and SureSpin 630 rotor (Thermo Scientific) at 20,000 rpm for 1.5 hours at 18°C. Viral stock titers were obtained by a limiting dilution assay (TCID_50_) assay on MRC-5s or ARPE19s. For titers from time courses, ARPE19 epithelial cells were plated onto 6-well dishes at a density of approximately 300,000-500,000 cells/well and allowed to grow until confluent and growth arrested for at least two days for cell cycle synchronization. Cells were infected at the indicated MOI using an approximation of 1.2 x 10^6^ cells per confluent well. Output virus was titered using an infectious units assay. Confluent ARPE19 cells (ATCC) were infected with viral inoculum collected at the indicated timepoints, incubated at 37°C with rocking for 2 h, and media was replaced. Titers were collected at 48 hpi by fixing with methanol. Cells were labeled with mouse anti-IE1 (1:500) primary antibody for 1 h at room temperature and secondary antibody Alexa fluor-488 goat anti-mouse (1:1000) for 1 h at room temperature. IE1-positive cells per well were quantified, divided by volume of viral inoculum, and titers reported as IU/ml.

### Nucleic Acid Analysis

Cellular and viral DNA levels were determined using quantitative polymerase chain reaction (qPCR) analysis. After collection, DNA was extracted using the DNeasy Blood & Tissue Kit (Qiagen) and resuspended in 100 μl. qPCR was performed using primers for HCMV UL123 and cellular CDKN1A. Primer sequences can be found below. Quantification was achieved by using Power SYBR Green PCR Master Mix (ThermoFisher Scientific) and QuantStudio 6 Flex real-time PCR System (ThermoFisher Scientific). qPCR began with an initial denaturation phase at 95°C for 20 s. DNA synthesis was performed for 40 cycles beginning with a melting phase at 95°C for 1 s and primer annealing and elongation phases at 60°C for 20 s. An absolute standard curve was constructed using plasmids pLVX-IE1-HA generously provided by Dr. Michael Nevels (43) and pLVX-CDKN1A-HA (16). Plasmid concentration was measured and copies/μl were determined by using a DNA copy number calculator (ThermoFischer Scientific). Absolute quantification for one biological replicate was obtained from the average quantities from at least two technical replicates. The three biological replicates were averaged, and the SD was obtained. The primer sequences for HCMV UL123 were 5′-GCCTTCCCTAAGACCACCAAT-3′ and 5′-ATTTTCTGGGCATAAGCCATAATC-3′ and for cellular CDKN1A were 5-GCGACTGTGATGCGCTAAT-3’ and 5’GTGGTGTCTCGGTGACAAAG-3’. Input vDNA decay analyses as completed by using 1×10^6^ growth arrested MRC5s were infected at intended vDNA_in,0_ 0.9, 11, and 14 genomes/cell with a different viral stock for each of two biological replicates. A MOI versus vDNA_in,0_ calibration curve was generated for each viral stock and the MOI needed to obtain the intended vDNA_in,0_ was back calculated from the calibration curve. After the 2 h infection protocol described in (16), the media was replaced with either normal DMEM or DMEM with 10 mM ganciclovir (GCV) and 30 mg/mL PFA. Media was replaced every 24 h.

### Protein Analysis

Protein levels were analyzed via immunoblotting. Cells were resuspended in Protease Inhibitor Cocktail (Sigma) and lysis buffer (1.5% SDS, 10 mM NaCl, 50 mM Tris Cl (pH 7.2), 1 mM EDTA) and lysed by sonication. Total protein concentration was determined via Pierce BCA Protein Assay Kit (Thermo Fisher Scientific). Proteins were separated by adding 20 μg of protein lysate to 4-20% gradient Mini Protean TGX Stain Free Pre-Cast Gels (Bio-Rad Laboratories). Proteins were then transferred to a nitrocellulose membrane (Cytiva) using the semi-dry Trans-Blot Turbo Transfer system (Bio-Rad). The membrane was blocked in 5% milk in TBS-T (tris-buffered saline, 0.1% Tween 20) for 1 h at room temperature, incubated in primary antibody overnight at 4 °C with 5% milk in TBS-T. Blots were washed in TBS-T, incubated in either fluorescent or horseradish peroxidase (HRP)-conjugated secondary antibody for 30 min or 1 h, respectively, in 5% milk at room temperature and imaged using ChemiDoc MP Imaging System (Bio-Rad). Antibodies used for immunoblotting can be found below. Sample lanes were normalized to the total protein in the undiluted internal standard lane and then normalized to the antibody-specific signal present in the undiluted internal standard lane present in all six blots across MOIs. Each replicate was normalized such that the 96 hpi time point had a standard deviation of 0, which was achieved by finding a normalization factor that could be multiplied by the value of the replicate at 96 hpi so as the value of the replicate was equal to the average. This normalization factor was then applied to each time point in the replicate. This process was performed for each replicate independently. The values from each replicate were averaged and the standard deviation (SD) was obtained. Primary antibodies used for immunoblots were mouse anti-UL44 (ICP36; 1:10,000; Virusys) and mouse anti-pp28 (1:1,000) HCMV antibodies were generously provided by Dr. Thomas Shenk (44). Secondary antibodies used for immunoblots were StarBright Blue 520 goat anti-mouse (IgG; 1:10,000; Bio-Rad) and Horseradish peroxidase conjugated-goat anti-mouse (IgG; 1:10,000; Jackson Immuno Research).

### Incucyte Live Cell Analysis and Confocal Microscopy

Infected MRC-5 or ARPE19 cells were place into Incucyte S3 (Sartorius) at 2 hpi. Images were taken every 2 h beginning at 3 hpi at 20x magnification with a green fluorescent filter (460 nm, IE2-2A-eGFP), red fluorescent filter (585 nm, pp28-mCherry), and phase contrast image with 16 images taken per well in each replicate. Scans range from 3 to 97 hpi. Images were analyzed and quantified using the Incucyte software. For confocal microscopy analysis, ARPE19 epithelial cells were plated on six-well dishes containing glass coverslips. Cells were serum starved with 0.5% FBS containing medium for 24 h for growth arrest. Once synchronized, cells were infected with TB40_Epi_ or TB40_Fb_ at an MOI of 0.5 IU/cell, washed at 2 hpi, then fresh medium was applied. At 96 hpi cells were fixed with 4% paraformaldehyde for 20 min and permeabilized in 0.1% Triton X-100 for 15 min, followed by blocking with 5% normal goat serum (Millipore-Sigma) in 3% BSA for 1 h. IE2 anti-mouse antibody (Clone 3A9, 1:200) generously provided by Tom Shenk was diluted in 5% normal goat serum and 3% BSA overnight at 4° C. Cells were washed with PBS-T and incubated for 1 h with secondary antibody conjugated to Alexa Fluor 647 (1:1000) and diluted in 5% normal goat serum in 3% BSA at room temperature. Cells were stained with Hoechst (1:1000) for 10 min at room temperature in the dark. Coverslips were placed on glass slides and mounted with ProLong Glass Antifade Mountant (Thermo Fisher Scientific). Images were acquired on a Nikon Ti2 CSU W1 spinning disk confocal microscope, and NIS elements was used for imaging.

### Model Parameterization and Simulation

All computational modeling studies were conducted in MATLAB R2021b (MathWorks).

#### Reduced Empirical Model of vDNA Replication

The empirical model of vDNA replication from (16) accounts for the major species of vDNA during lytic replication. In this model, vDNA_tot_ is defined to be the sum of input vDNA (vDNA_in_) and replicated vDNA (vDNA_rep_). Given that different species of vDNA were indistinguishable in our qPCR assay, the vDNA_tot_ variable was fit to qPCR data using a strategy similar to that proposed in (16). Briefly, the built-in MATLAB constrained optimization function “fmincon” was used to simultaneously estimate parameters *k_d_, V_max_, t_50_*, and *nH* for all experimentally quantified vDNA_in,0_ for each infection condition by fitting the model proposed in (16) to the mean qPCR data in **Fig. 1E**. In addition, individual biological replicates were fit using the same procedure and individual parameter estimates were obtained. These parameters were then plotted in **Fig. 2B** and analyzed using a Kruskall-Wallace test with Dunnett’s correction for multiple comparisons (Prism, GraphWorks). Importantly, for this and subsequent modeling analyses given the variability in vDNA at 2 hpi between infection conditions, vDNA_in,0_ was back calculated from the estimated *k_d_* parameter and the vDNA at 2 hpi to obtain a more accurate vDNA_in,0_ representing vDNA at time, t = 0 hpi.

#### Mechanistic Model of Late Lytic Replication

The mechanistic model of late lytic replication is described in (16). Briefly, the model schematic described in **Fig. 5A** was created by synthesizing known mechanisms of viral protein synthesis, capsid formation, nuclear egress, and virion production. From this schematic and using the laws of mass balance, mass action, and Michaelis-Menten kinetics, the coupled ordinary differential equations (ODEs) in **Fig. 5B** were generated. The model selected by the authors in (16) and presented in this manuscript was, in fact, selected from a series of six competing models by balancing parsimony with the addition of novel kinetic mechanisms. For further information on model selection, please refer to (16). The ODE-based mechanistic model of late HCMV lytic replication was then fit using a simplified version of the pseudo-Monte Carlo parameter estimation approach described in (16). Parameters found in (16) were rescaled by hand to coarsely fit the re-analyzed immunoblot data presented in this manuscript and from (16) (see **Methods, Protein Analysis**). Operating under the assumption that there is no difference in parameters between experimental conditions (i.e., null hypothesis), these hand-tuned initial guesses were then used as initial guesses for subsequent pseudo-Monte Carlo parameter estimation. The process of pseudo-Monte Carlo parameter estimation is described in detail in (16). Briefly, hand-tuned initial guesses were optimized using 1000 iterations of the built-in MATLAB constrained optimization algorithm “fmincon.” Optimal parameters were selected by examining histograms of parameter estimates for each of the 1000 iterations (see **Fig. 7**). From these histograms, we found the most frequently estimated value for each parameter and eliminated outlier values within +/− 20% of the most frequently estimated value. The remaining estimated values for each parameter were then averaged and used in simulations. For fitting, parameter estimates were constrained between 0.1 and 10 times of the initial guess values and k_d_ were fixed as described in (16). K_m_ parameters were also fixed as they broadly represent intrinsic protein affinities for substrate determined by amino acid sequence and were hypothesized to be less likely to change between infection condition. Conversely, activity parameters such as the k_s_ constants are related to factors both intrinsic and extrinsic to the protein itself as these constants are generally the product of two quantities, the intrinsic catalysis rate and the concentration of active protein (45). We fit the ODEs to the TB40_Fb_ infection of fibroblast data from (16) using the procedure described above to generate values for the K_m_ parameters. As the values for the K_m_ parameters were published previously in (16), we were able to compare our estimated to those previously established. Each of the K_m_ values were then fixed to the value estimated for TB40_Fb_ infection of fibroblast for subsequent parameter estimation of each infection condition.

## ACKNOWLEDGEMENTS

The authors are thankful to the Terhune, Dash, and Hudson Lab members for their input during the development of this work. We thank Drs. Laura Hertel and Mike McVoy for discussions on virus stock nomenclature. We thank Drs. Eain Murphy, Michael Nevels, and Tom Shenk for providing recombinant HCMV, UL123 plasmid, and antibodies against HCMV proteins, respectively. Schematics were generated using BioRender.com.

This work was supported by the National Institutes of Allergy and Infectious Disease Division of the National Institutes of Health under award number F30AI179084 to CEM, R21AI149039 to SST and RKD, and R01AI083281 to SST. The content is solely the responsibility of the authors and does not necessarily represent the official views of the National Institutes of Health. The funders had no role in study design, data collection and analysis, decision to publish, or preparation of the manuscript.

## Author Contribution Statement

R.L.M. and S.S.T. designed *in vitro* experiments; R.L.M., S.R.R, and M.L.S. performed *in vitro* experiments; C.E.M. and R.K.D. developed and tested *in silico* computational models and performed simulations; R.L.M., C.E.M., S.R.R., M.L.S., R.K.D., and S.S.T. analyzed the data; and R.L.M., C.E.M., S.R.R., M.L.S., R.K.D., S.S.T. wrote and edited the manuscript and approved the final submission.

## Classification

Microbiology; Virology; Systems Biology; Computational Biology; Modeling Biological Systems.

